# Seasonal microbiome community dynamics in the massive coral *Porites lobata* impacted by sedimentation

**DOI:** 10.64898/2026.05.08.723747

**Authors:** Justin T. Berg, James E. Fifer, Sarah W. Davies, Bastian Bentlage

## Abstract

Near-shore coral reefs in southern Guam (Mariana Islands) experience severe sedimentation, in particular during the wet season when rainfall and erosion are high. We sampled fragments of the reef-forming coral *Porites lobata* from opposite ends of a sedimentation gradient in Fouha Bay, southern Guam, during dry and wet seasons. Using DNA metabarcoding, we characterized the diversity and composition of *P. lobata*-associated Symbiodiniaceae and bacterial microbiome communities. As in many species of *Porites*, Symbiodiniaceae communities of *P. lobata* were dominated by variants of *Cladocopium* C15 with sites showing differences in Symbiodiniaceae communities attributable to variation in these *Cladocopium* C15 variants. Bacterial microbiomes of *P. lobata* were dominated by *Endozoicomonadaceae*, a family of putative coral bacterial endosymbionts involved in nutrient cycling. Site and seasonal differences in bacterial diversity and community composition were apparent. In close proximity to the mouth of the river draining into Fouha Bay, bacterial diversity was highest during the wet season when sedimentation is generally severe. Microbiome reorganization in response to sedimentation may explain this result, but we also found overrepresentation of bacteria associated with terrestrial origin close to the river mouth and/or during the wet season. Together these patterns highlight that coral Symbiodiniaceae and bacterial communities are both spatially and temporally structured in this disturbed system.

**IMPORTANCE:** This study provides a time series dataset of coral-associated microorganisms, including dinoflagellate algae and bacteria, from a tropical bay impacted by sedimentation that results from upstream erosion of disturbed soils. Characterizing temporal patterns of coral-associated microbes provides insights into the dynamic nature of these communities. While microbiome variability across sites and seasons may be a result of acclimatization to different environmental conditions, we identified bacterial groups of putative terrestrial origin in sampled coral microbiomes that may have been exported from eroded soils to the near-shore reef. Considering that disturbed soils act as hotspots for the proliferation of potentially harmful substances, such as antimicrobial resistance genes, understanding microbial community connections at the marine-freshwater-terrestrial interface is an important step toward evaluating environmental impacts across connected ecosystems from ridge to reef.

## INTRODUCTION

Erosion and resulting sedimentation represent major threats to nearshore marine ecosystem function for islands such as Guam (Abraham et al., 2004; Richmond, 1993). Increasing sedimentation due to poor upstream soil management and land use practices is a major cause for declining coral community species diversity and abundance near river mouths (Macdonald et al., 1997; Ramos-Scharrón & MacDonald, 2005; Rongo, 2004). In southern Guam, challenges to land management led to severe sedimentation on nearshore reefs, with Fouha Bay representing one of the best documented cases of these impacts (Wolanski et al., 2004). Fouha Bay’s reef comprised 155 coral species close to fifty years ago (Birkeland and Randall, 1981). Road construction in the 1980s led to major erosion and nearshore sedimentation which led to a reduction of coral diversity to less than 92 species present some two decades later (Rongo, 2004). Wildland arson continues to disturb vegetation in the Fouha Bay watershed, leading to the formation of badlands and high levels of erosion (Porter et al., 2005; Minton, 2006). On average, Fouha Bay experiences ten major sedimentation events annually, but flushing through storm-driven swells only occurs two to five times annually, causing retention of deposited sediments (Wolanski et al., 2003).

Sedimentation in Fouha Bay decreases with increasing distance from the mouth of the La Sa Fua River (Rongo, 2004; Wolanski et al., 2003). Inversely, coral species richness increases with distance from the river mouth with the reef-building *Porites lobata* and encrusting *Leptastrea purpurea* representing the only coral species that persist under a regime of chronic, severe sedimentation in the inner bay (Minton, 2022). Severe sedimentation smothers coral tissues, leading to necrosis (Erftemeijer et al., 2012; Tuttle and Donahue, 2022), negatively impacts coral metabolism (Fabricius, 2005; Bollati et al., 2022), reduces reproductive output (Ricardo et al., 2015), and inhibits photosynthesis of coral-associated Symbiodiniaceae photo-endosymbionts (Rushmore et al., 2021). In addition, sedimentation can alter coral-associated microbial communities, which may, for example, lead to diseases (Bruno et al., 2007; Mouchka et al., 2010). Coral bacterial endosymbionts are integral to coral holobiont function and homeostasis (Thurber et al., 2020) and may mitigate adverse environmental conditions (Reshef et al., 2006; Rosenberg et al., 2007). Among the key coral endosymbionts are bacteria of the family *Endozoicomonadaceae* (Gammaproteobacteria: Oceanospirillales) (Pernice et al., 2020), which are diverse and prevalent across the Pacific Ocean (Galand et al., 2023). *Endozoicomonadaceae* are thought to be positively associated with coral health, and diseased corals are often characterized by reduced abundances of *Endozoicomonadaceae* (Bayer et al., 2013; Glasl et al., 2016; Meyer et al., 2014; Morrow et al., 2017; Neave et al., 2016; Vezzulli et al., 2013). Some strains of *Endozoicomonadaceae* have been shown to produce antimicrobial compounds (Ritchie, 2006; Rua et al., 2014), which could act as a biological control for pathogens and prevent negative impacts of pathogens present in the environment, as may be expected when sedimentation is high.

Fifer et al. (2022) showed that *P. lobata* bacterial microbiomes in Fouha Bay were dominated by *Endozoicomonadaceae* with other less abundant bacterial taxa showing differential abundance along the sedimentation gradient. In addition, Symbiodiniaceae communities were dominated by *Cladocopium* C15 strains, with no apparent structuring of ITS2 profiles (Fifer et al., 2022). In this contribution, we expand on Fifer et al.’s (2022) study that focused on a single timepoint. Inner and outer sites, characterized by severe and moderate sedimentation, were sampled monthly in Fouha Bay to identify potential seasonal dynamics of *P. lobata*-associated Symbiodiniaceae and bacterial microbiome communities using a DNA metabarcoding approach. Guam’s climate is characterized by distinct dry and wet seasons (Gingerich, 2003). As such, sedimentation is higher during the wet season that sees high precipitation compared to the dry season (Lock et al., 2024). We anticipated that proximity to the river mouth and season affect endosymbiont communities of *P. lobata*. Shuffling of Symbiodiniaceae endosymbionts has been proposed as a mechanism for corals to respond to environmental stress (Cunning et al., 2015; Lan et al., 2025), but we found that Symbiodiniaceae remained stable across seasons but differed between sites. By contrast, bacterial microbiomes showed diversity and compositional differences between sites and seasons.

## MATERIALS AND METHODS

### Environmental data collection

A YSI 6-Series Multiparameter Water Quality Sonde (Yellow Springs, Ohio) was deployed at two sites (Fig. 1) to measure water temperature and conductivity during two months of the wet season (September and November) and two months of the dry season (January and March). River discharge data for the La Sa Fua river were obtained from the U.S. Geological Survey (2019-2020 data from monitoring station 16809600) for the same time period.

**Fig. 1.**
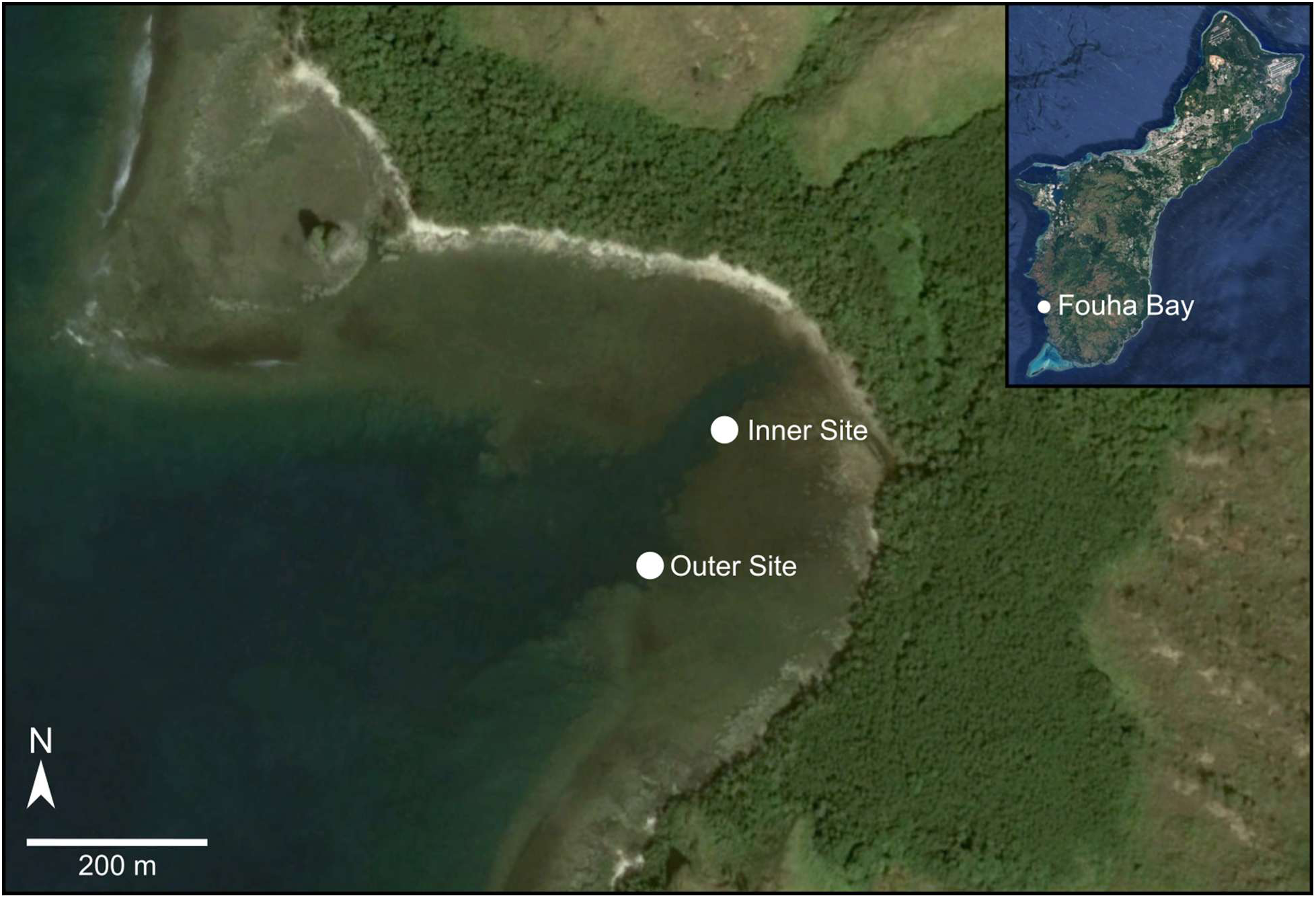
Inner and Outer sampling sites in Fouha Bay; location of Fouha Bay in southern Guam shown in upper right hand corner inset. Maps created using Google Earth Pro with images from Maxar Technologies and Landsat/Copernicus.

### Coral sample collection

Eight colonies of *Porites lobata* growing near the river mouth in Fouha Bay (inner site; Fig. 1) and eight colonies growing near the mouth of the bay (outer site; Fig. 1) were tagged in September 2019. Colonies were sampled repeatedly eight times at roughly 1-month intervals from September 2019 to May 2020 spanning a 3-month period during the wet season and a 5-month period during the dry season. In total, 126 samples were collected from the center of coral colonies with a hammer and chisel, placed individually in a pre-labeled Whirl-Pack (Consolidate Plastics, Stow, OH), and immediately frozen in liquid nitrogen, followed by storage at -80**°**C until DNA extraction.

### ITS2 and 16S metabarcoding

DNA was extracted using the DNeasy PowerSoil Kit (Qiagen, Hilden, Germany) in conjunction with a QIAcube Connect liquid handling robot (Qiagen, Hilden, Germany) following the manufacturer’s protocol. DNA was quantified using a Quibit fluorometer (Qubit, Carlsbad, CA) prior to sending DNA aliquots to CD Genomics (Long Island, NY) for PCR amplification and metabarcoding on an Illumina NovaSeq 6000 sequencer (Illumina, San Diego, CA).

Symbiodiniaceae ITS2 was amplified using the protocol of McDevitt-Irwin et al. (2017) and the v4 hypervariable region of bacterial 16S was amplified using primers 515f (Parada et al., 2016) and 806r (Walters et al., 2015).

### Symbiodinaceae ITS2 data analysis

Raw paired-end ITS2 reads were uploaded to SymPortal (Hume et al., 2019) for processing and taxonomic assignment. Only those samples that passed quality filtering for bacterial 16S data analysis (see below) were retained for Symbiodiniaceae community analysis.Because ITS2 is present in multiple copies within Symbiodiniaceae genomes and copy number varies among taxa, diversity metrics are often interpreted cautiously for Symbiodiniaceae community datasets (Davies et al., 2023). However, because the majority of Symbiodiniaceae sequence variants recovered in this study belonged to *Cladocopium* C15, alpha diversity metrics were explored here as a within-lineage assessment of Symbiodiniaceae variability. Observed diversity (number of unique features), Shannon diversity, and evenness were calculated to compare alpha diversity across sampling groups.

Initially, linear mixed-effects models including coral colony as a random effect were evaluated for bacterial alpha-diversity metrics; however, singular model fits indicated negligible variance associated with the random-effect structure. Because repeated observations were also incomplete due to sample loss, subsequent analyses were conducted using fixed-effects analysis of variance (ANOVA). ANOVA was used to assess the effects of site, season, and their interaction on ITS2 alpha diversity metrics. Tukey’s Honest Significant Difference (HSD) tests were used for post hoc comparisons following significant ANOVA results. The contribution of site, season and their interaction to the variation in Symbiodiniaceae beta diversity patterns was evaluated using permutation analysis of variance (PERMANOVA) through the adonis function using default settings (9999 permutations) in the vegan R package (Oksanen et al., 2001); by default, the adonis function uses Bray-Curtis distances. An analysis of similarity (ANOSIM) with 9999 permutations was used to test for significant group differences.

### Bacterial 16S data analysis

Following demultiplexing and removal of primer and barcode sequences by CD Genomics using their in-house pipeline, including merging of paired reads using the Fast Length Adjustment of SHort reads (FLASH) software (Magoč & Salzburg, 2011). Contigs were imported into QIIME2 (Boylen et al., 2019), truncated to a maximum length of 253bp and amplicon sequence variants (ASV) were inferred using DADA2 (Callahan et al., 2016). ASVs were assigned taxonomies using the Greengenes classifier version 13_8 (DeSantis et al., 2006). ASVs inferred as mitochondrial or chloroplast origin were removed from the dataset. In addition, rare ASVs occurring in <5% of samples or with a frequency <0.01% were removed prior to further analysis. ASVs remaining after filtering were aligned using the QIIME MAFFT plug-in (Katoh & Standley, 2013) followed by masking of highly variable alignment positions. To allow for the calculation of phylogenetic distances between ASVs, the maximum likelihood tree for the ASV alignment was inferred using the QIIME2 implementation of IQ-TREE (Minh et al., 2020); the phylogeny was rooted using the midpoint rooting method.

To ensure equal sampling depth across all 112 samples remaining after data processing and quality filtering, ASVs were rarefied to a depth of 80,000 reads per sample. Shannon diversity and evenness were calculated using the QIIME2 core metrics plug-in. Initially, linear mixed-effects models including coral colony as a random effect were evaluated to account for repeated sampling structure. However, singular model fits indicated negligible variance associated with the random effect for alpha-diversity metrics. In addition, repeated observations were incomplete due to sample loss across time points. Therefore, subsequent analyses were conducted using fixed-effects ANOVA. ANOVA and Tukey’s HSD tests were used to evaluate the effects of site, season, and their interaction on bacterial alpha diversity. Phylogenetic distances were used to calculate unweighted UniFrac distances (Lozupone & Knight, 2006) for beta diversity analysis; unweighted rather than weighted UniFrac distances were used to reduce the influence of highly abundant, dominant ASVs on the analysis. ANOSIM, as implemented in QIIME2, was used to test if groups (inner dry, inner wet, outer dry, and outer wet) were significantly different. A PERMANOVA using the QIIME2 Adonis (Simpson et al., 2022) plug-in with 9999 permutations was used to estimate how much of the variation in unweighted UniFrac distances is explained by site, season, and their interaction term. Analysis of compositions of microbiomes with bias correction (ANCOM-BC; Lin & Peddada, 2020), as implemented in QIIME2, was used to identify differentially abundant taxa. A Pearson Correlation Analysis was performed using the Shannon Diversity scores for the Symbiodiniaceae and bacterial communities to better understand possible correlations between these symbiotic communities.

## RESULTS

### Environmental Data

Temperatures ranged from ∼26**°**C to ∼31.5**°**C and were consistent between inner and outer sites; water temperatures dropped by ∼2**°**C between the tail-end of the wet season in November and the dry season in January (Fig. 2A). Conductivity ranged from ∼55 to ∼40 mS/cm with consistent differences between inner and outer sites (Fig. 2B). During the wet season in September and November, conductivity was lower in the inner site than the outer site. This is consistent with high monthly river discharge and precipitation during the same time period (Figs. 2C & D). By contrast, during the dry season this river discharge and precipitation decreased (Figs. 2C & D) while conductivity in the inner site increased (Fig 2B).

**Fig. 2.**
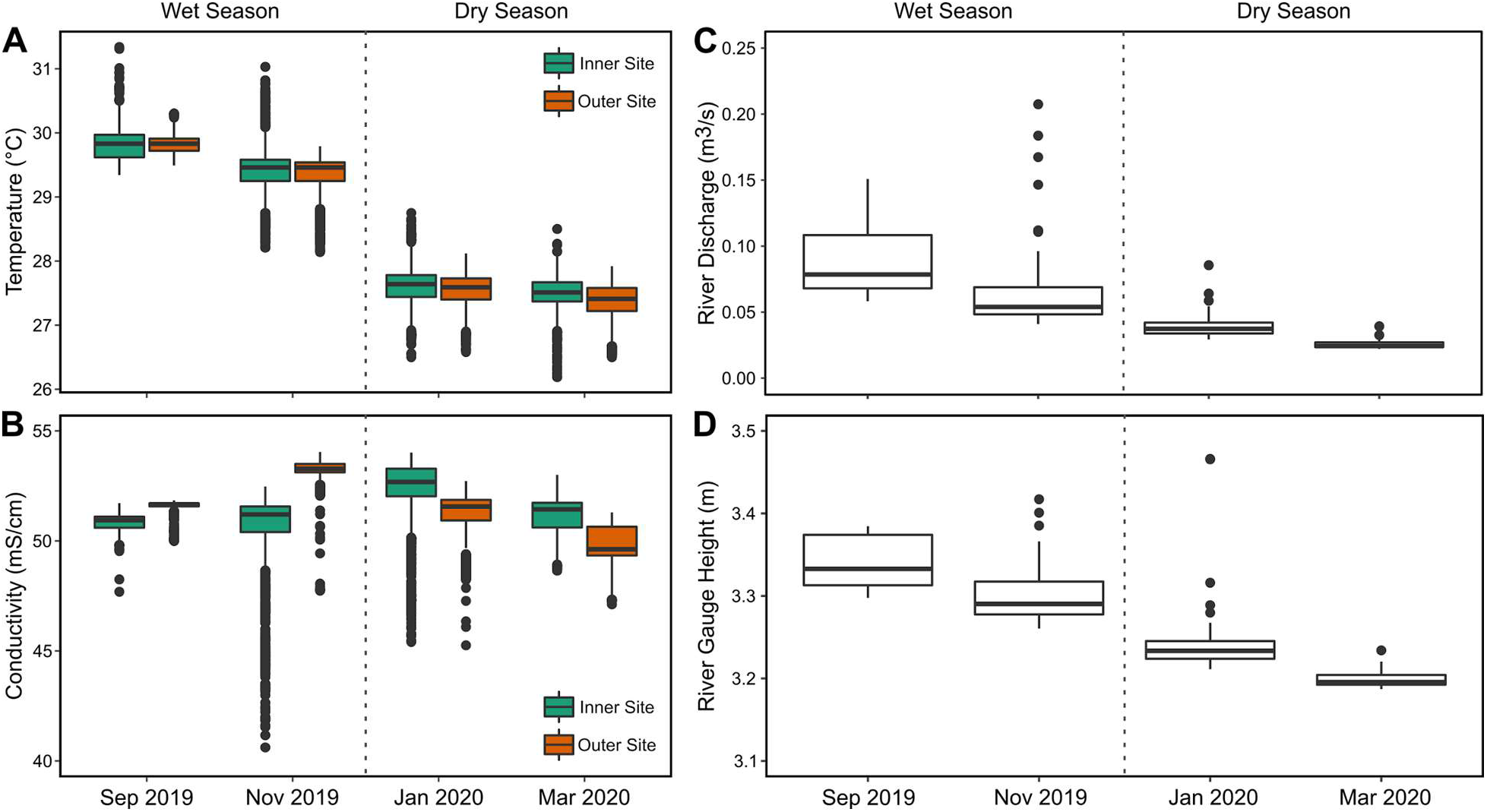
Environmental data collected during the wet (September and November 2019) and dry (January and March 2020) seasons. Water temperature (A) and conductivity (B) were recorded in Fouha Bay using YSI data loggers. Discharge (C) and gauge height (D) for the La Sa Fua river draining into Fouha Bay were obtained from the United States Geological Survey.

### DNA sequencing and data deposition

Sequence data generated by this project were deposited in NCBI’s GenBank under BioProject PRJNA1313140. 126 samples were successfully amplified using Symbiodiniaceae specific ITS2 primers and yielded from 67,503 to 92,707 (average of 84,371) paired-end sequences per sample that were deposited in the SRA under accession numbers SRR35197709–SRR35197834. 115 samples were successfully amplified using bacterial 16S primers and yielded from 91,581 to 233,640 (average of 184,849) paired-end sequences per sample that were deposited in the short read archive (SRA) under accession numbers SRR35197587–SRR35197701.

### Symbiodiniaceae diversity and community composition

ITS2 metabarcoding showed that *Porites lobata* colonies in Fouha Bay were dominated by variants of *Cladocopium* C15 across sites and seasons (Fig. 3; Supplementary Fig. 1) with smaller contributions of other *Cladocopium* clades and *Gerakladium* G3. Symbiodiniaceae alpha diversity differed significantly between sites in observed features (p < 0.001), evenness (p < 0.001), and Shannon diversity (p < 0.05) (Fig. 4) (Supplementary Tables 1-3). Consistent with the results for alpha diversity comparisons, PERMANOVA identified site as the major factor explaining Symbiodiniaceae community composition (p < 0.001) while season and the interaction term between site and season were not significant contributors to the variation in beta diversity (Table 1). Consequently, a high degree of dissimilarity between Symbiodiniaceae communities was identified by ANOSIM across sites (R = 0.731, p = 0.0001, 9999 permutations).

**Fig. 3.**
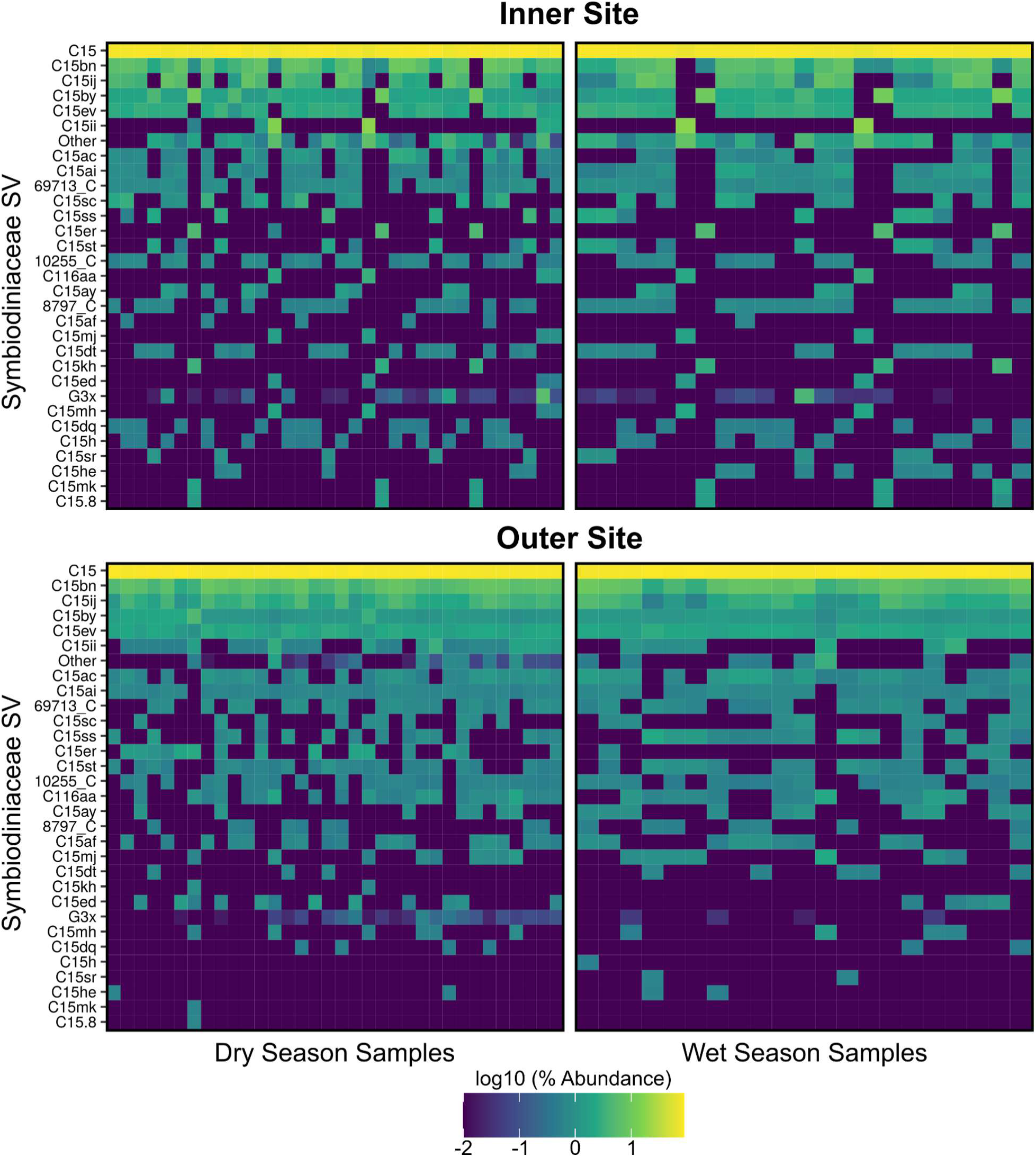
Symbiodiniaceae community composition during dry and wet seasons at inner and outer sites. Columns of the heat map represent individual samples; rows represent Symbiodiniaceae clades. Relative abundances were log-scaled using the common logarithm where 2 equals 100% (yellow) and -2 equals 0.01% (blue). Symbiodiniaceae clade abbreviations: C = *Cladocopium*; G = *Gerakladium*. SV = sequence variant.

**Fig. 4.**
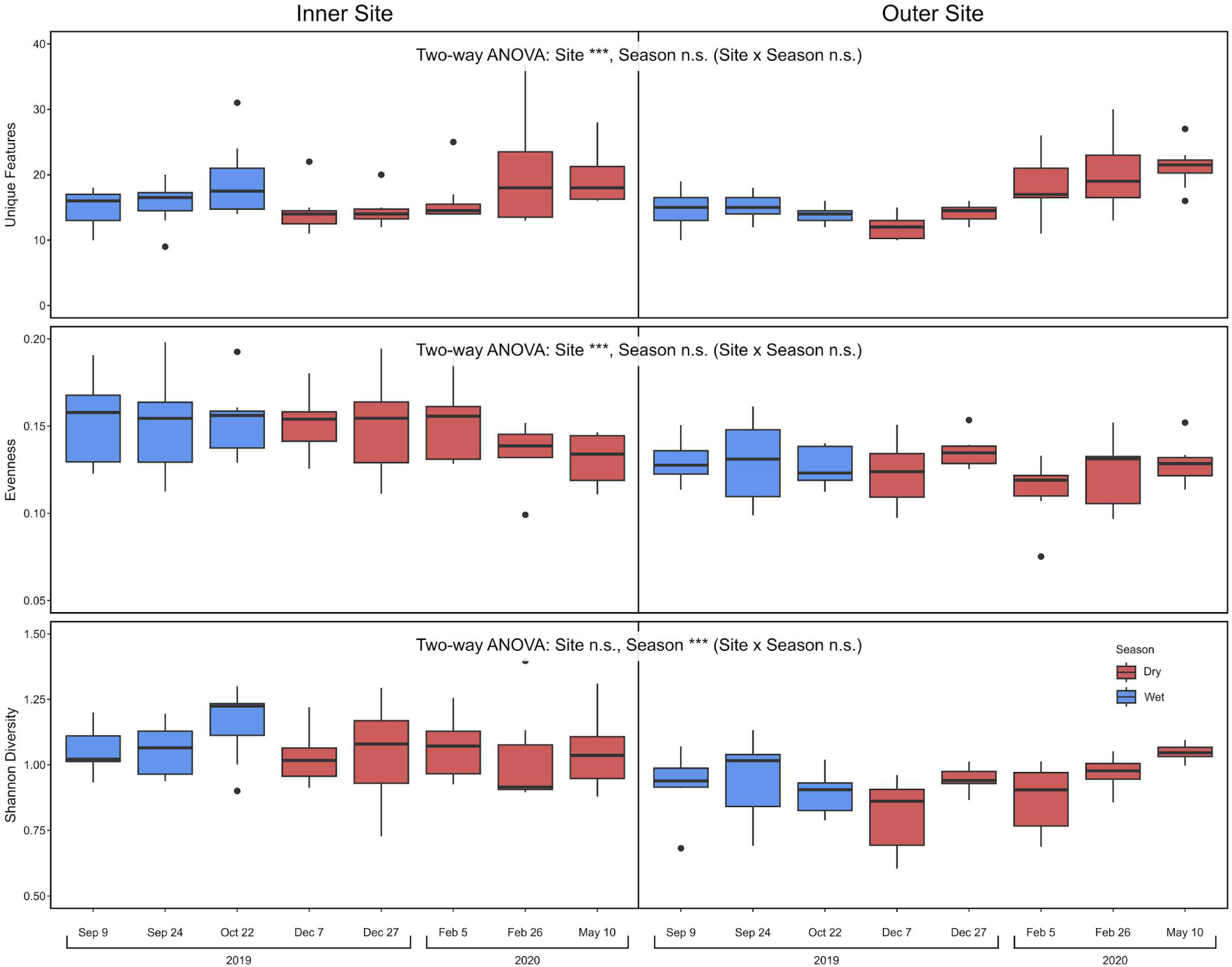
Symbiodiniaceae alpha diversity for *Porites lobata* samples collected at inner and outer sites in Fouha Bay, Guam during the dry (red) and wet (blue) seasons.

**Table 1.** PERMANOVA table of the contributions of site, season and their interaction term on Symbiodiniaceae community composition.

|  | **df** | **Sums of Squares** | **$R^2$** | **$F$** | **$p$** |
| --- | --- | --- | --- | --- | --- |
| **Site** | 1 | 1.40 | 0.5 | 107.5 | 0.001 |
| **Season** | 1 | 0.01 | 0.004 | 0.77 | 0.484 |
| **Site:Season** | 1 | 0.008 | 0.003 | 0.64 | 0.569 |
| **Residuals** | 108 | 1.40 | 0.498 |  |  |
| **Total** | 111 | 2.82 | 1.00 |  |  |

### Bacterial microbiome diversity and community composition

Alpha diversity of *P. lobata* bacterial microbiomes (Fig. 5) differed significantly between sites, seasons, and the interaction term of site and season (ANOVA and HSD, p<0.001) (Supplementary Tables 4-6). The bacterial HSD results indicated significant differences between individual site/season groups (Inner Wet = IW, Inner Dry = ID, Outer Wet = OW, Outer Dry = OD). Significant differences were found in observed features (IW:ID, OD:IW, and OW:IW), evenness (IW:ID, OD:IW, and OW:IW), and Shannon diversity (IW:ID, OW:ID, and OW:IW). The highest bacterial diversity was observed for the inner wet season (IW) group across all alpha diversity metrics (Fig. 5). Overall, the inner site wet season group was the most diverse (Fig. 5) while outer site sample diversity was only elevated during September and early December, the likely result of heavy rainfalls shortly before sampling. In addition, bacterial Shannon diversity was mildly correlated with Symbiodiniaceae Shannon diversity (R=0.28; p<0.01; Supplementary Fig. 2).

**Fig. 5.**
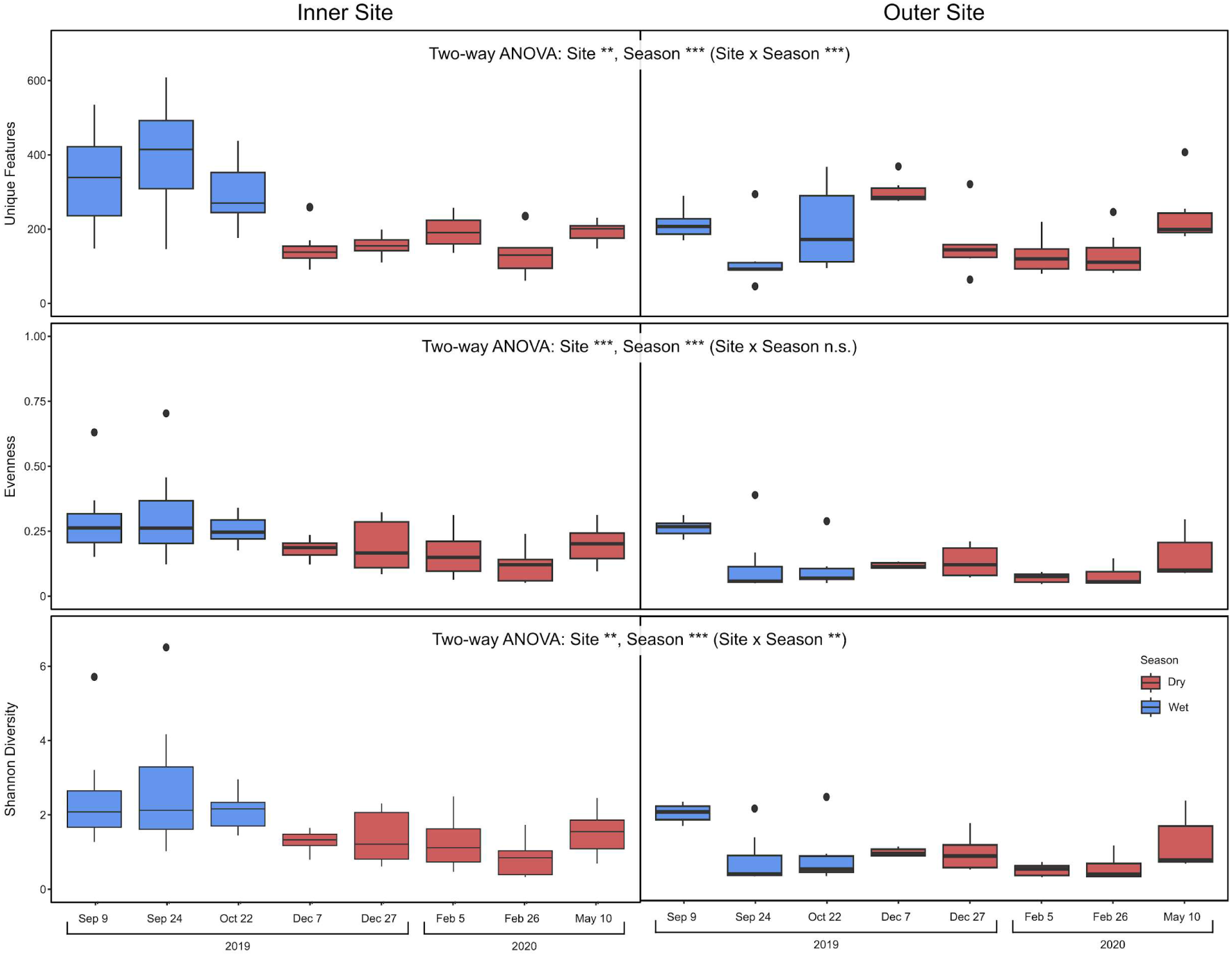
Bacterial alpha diversity for *Porites lobata* samples collected at inner and outer sites in Fouha Bay, Guam during the dry (red) and wet (blue) seasons.

Overall, the inner site, especially during the wet season, was characterized by relatively higher abundances of bacteria across taxonomic groups compared to the outer site, indicated by overall warmer colors in the heatmap comparison (Fig. 6; Supplementary Fig. 3). The *P. lobata* bacterial microbiome communities were dominated by *Endozoicomonadaceae* (Fig. 6; mean relative abundance of 82.5%) and *Oxalobacteraceae* (5.4%) across all samples. Inner site samples were dominated by *Endozoicomonadaceae*, *Oxalobacteraceae*, and *Spirochaetaceae* (mean relative abundance of 74.5.0%, 9.4%, and 1.5%, respectively) while outer site samples were dominated by *Endozoicomonadaceae* and *Vibrionaceae* (mean relative abundance at 90.8% and 1.2%, respectively); all remaining families had a relative mean abundance < 1%. Wet season samples were dominated by *Endozoicomonadaceae*, *Oxalobacteraceae*, and *Spirochaetaceae* (mean relative abundance of 78.7%, 1.5% and 1.6%) while dry season samples were dominated by *Endozoicomonadaceae* and *Oxalobacteraceae* (mean relative abundance at 85.0% and 7.8%); all remaining families had a relative mean abundance < 1%. The outer site dry season group was dominated by the family *Endozoicomonadaceae* (mean relative abundance at 93.7%). The outer site wet season group was dominated by the families *Endozoicomonadaceae*, *Oxalobacteraceae*, *Vibrionaceae* and *Enterobacteriaceae* (mean relative abundance at 86.3%, 1.4%, 2.0%, and 1.0%). The inner site dry season group was dominated by the families *Endozoicomonadaceae* and *Oxalobacteraceae* (mean relative abundance of 76.8% and 14.6%). The inner site wet season group was dominated by the families *Endozoicimonadaceae*, *Spirochaetaceae*, *Oxalobacteraceae*, *Lachnospiraceae*, and *Rhodospirillaceae* (71.7%, 3.0%, 1.7%, 1.4%, and 1.2%).

**Fig. 6.**
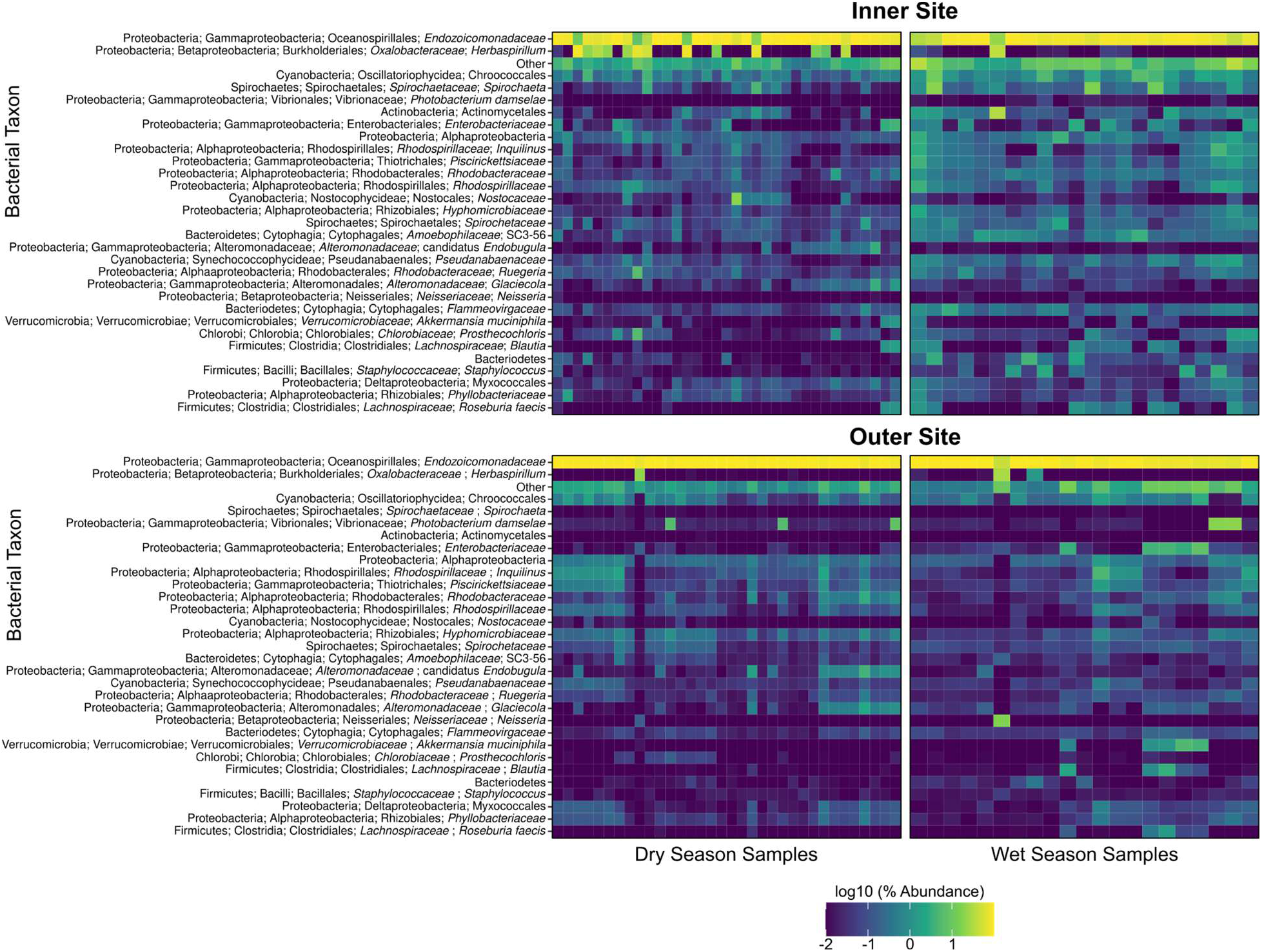
Relative abundance of the 30 most abundant bacterial ASVs in *Porites lobata* microbiomes sampled from an inner and outer site in Fouha Bay during the dry and wet season. Columns of the heat map represent individual samples; rows represent bacterial taxa inferred from ASVs. Relative abundance was log-scaled using the common logarithm where 2 equals 100% (yellow) and -2 equals 0.01% (blue).

Inner site bacterial microbiomes in the wet season showed limited overlap with the other three sample groups (inner site dry season, outer site wet and dry seasons; Fig. 7), a likely result of the comparatively high diversity of samples from the inner site wet season group (Fig. 5). As might be expected, inner site dry season samples overlapped with part of the distribution of inner site wet season samples. Outer site dry season samples were contained within the distribution of outer site wet season samples, a likely result of the dominance of *Endozoicomondaceae* and the limited amount of variation between seasons at the outer site. ANOSIM identified a significant degree of dissimilarity among site/season groups (p=0.0001, R=0.43, 9999 permutations), consistent with the differentiation of the inner site wet season group from the remainder of the samples (Fig. 7). The dominance of *Endozoicomonadaceae* (Fig. 6) likely explains the overall relatively low degree of differentiation indicated by the ANOSIM R value <0.5.

**Fig. 7.**
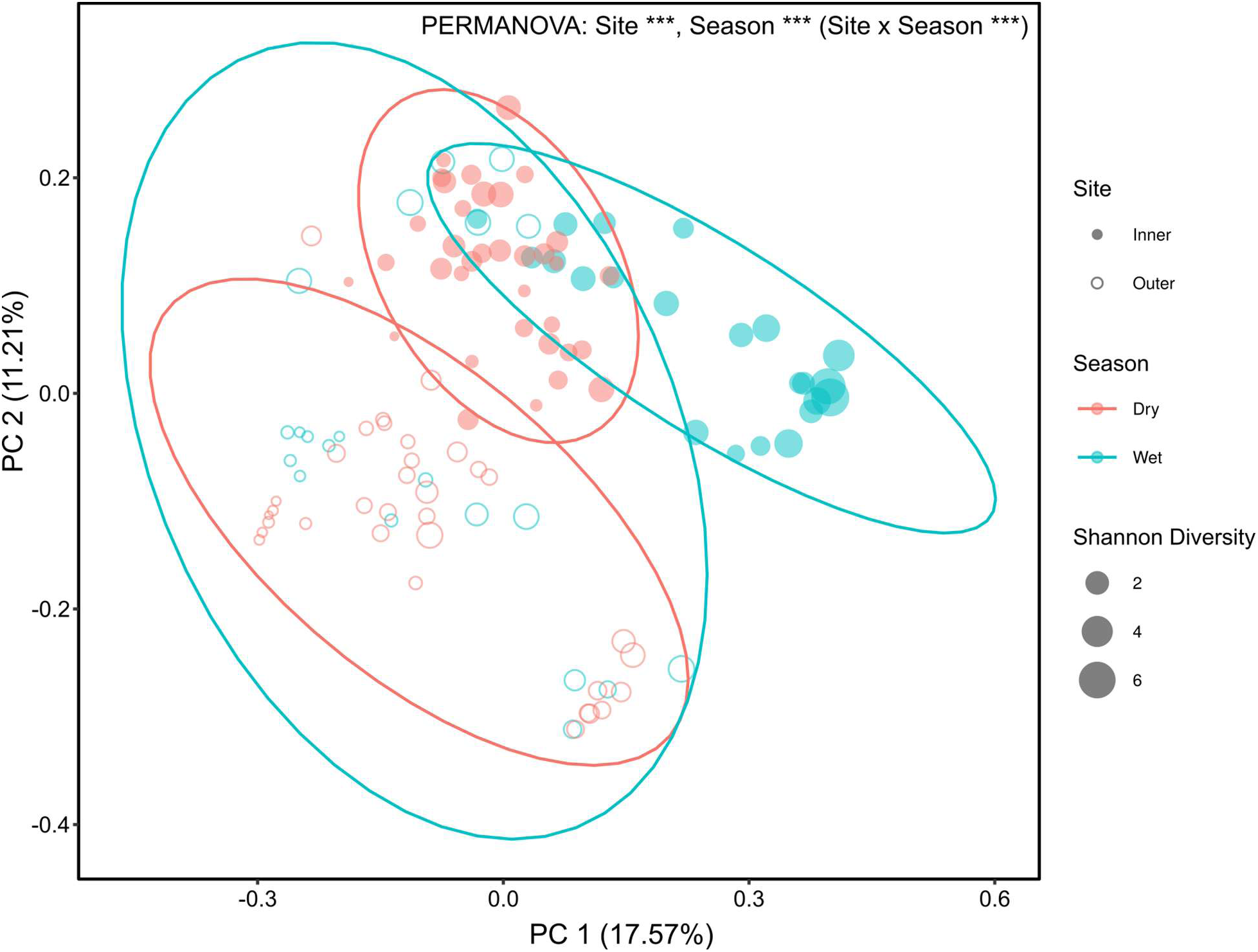
Unweighted unifrac PCoA of bacterial communities associated with *Porites lobata* from two sites in Fouha Bay, Guam during the wet and dry seasons. 95% confidence ellipses were calculated to highlight distributions of samples related to each by sampling site (inner *versus* outer site) and (dry *versus* wet season).

PERMANOVA identified site, season, and the combination of site/season as significant contributors to bacterial community diversity (Table 2). Considering that site (R^2^=0.11) and season (R^2^=0.05) contributed most to the observed variation compared to their interaction (site/season R^2^=0.03), ANCOM-BC was used to identify differentially abundant bacterial taxa that may explain the differences between sites and seasons. ANCOM-BC identified orders of Actinomycetota to be more abundant at the inner site compared to the outer site (Actinomycetales; Fig. 8A) and during the wet season compared to the dry season (Coriobacteriales; Fig. 8B). In addition, Deinococcales were more abundant at the inner site compared to the outer site (Fig. 8A); Firmicutes (Clostridiales) were more abundant during the wet season than the dry season (Fig. 8A).

**Fig. 8.**
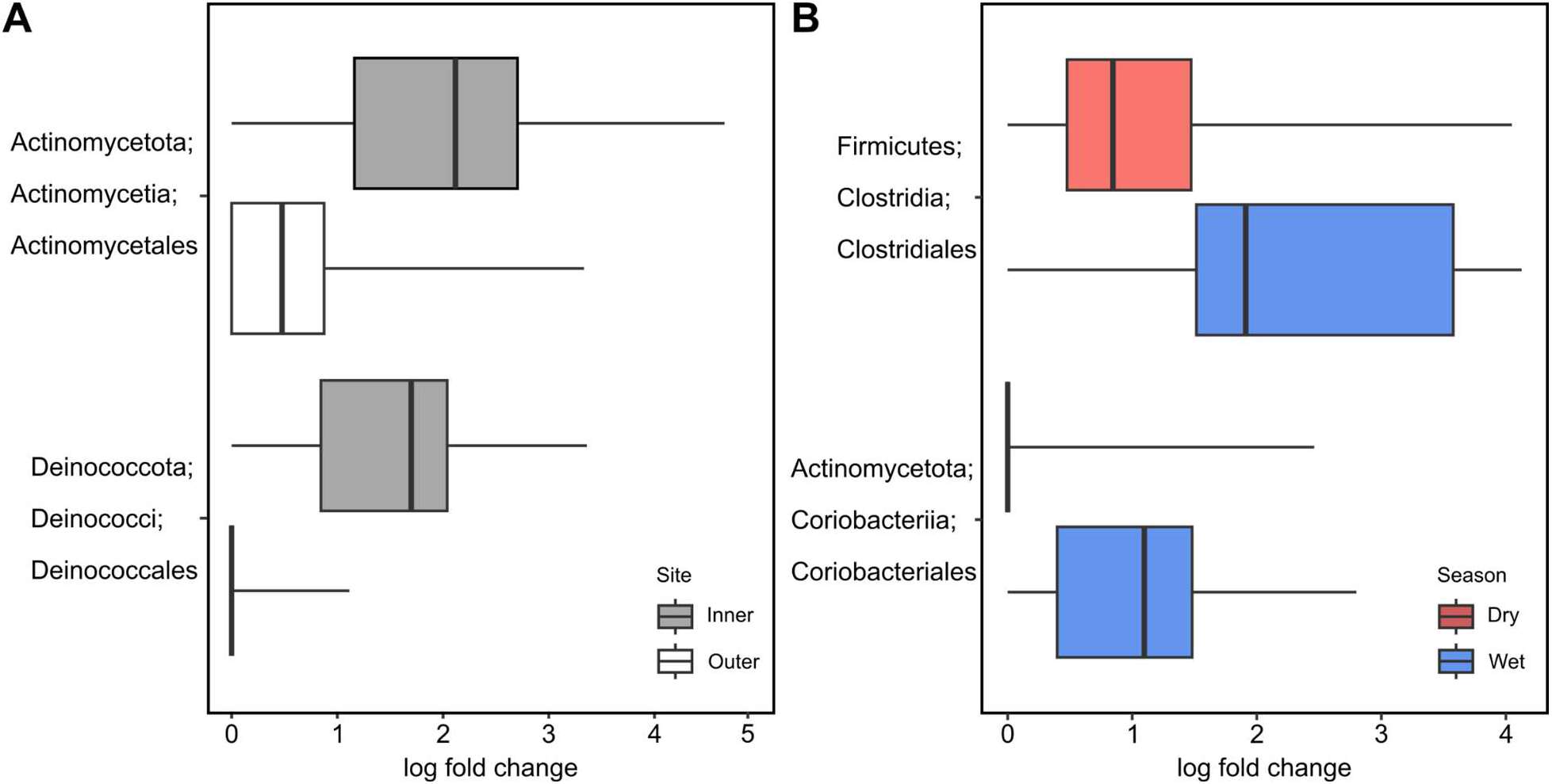
Significant log fold change differences in bacterial taxa at taxonomic level 4 identified by ANCOM-BC. Two bacterial groups were identified to be differentially abundant in comparisons of inner *versus* outer site (A) and dry *versus* wet seasons (B).

**Table 2.** PERMANOVA table of the contributions of site, season and their interaction term on bacterial microbiome composition.

| | df | Sums of Squares | $R^2$ | F | $p$ |
| --- | --- | --- | --- | --- | --- |
| Site | 1 | 2.09 | 0.11 | 14.23 | 0.001 |
| Season | 1 | 1.07 | 0.05 | 7.29 | 0.001 |
| Site:Season | 1 | 0.66 | 0.03 | 4.49 | 0.001 |
| Residuals | 108 | 15.87 | 0.81 |  |  |
| Total | 111 | 19.69 | 1 |  |  |

## DISCUSSION

### Environmental Differences in Fouha Bay

From September of 2019 to May of 2020, Fouha Bay showed stark differences in environmental parameters and microbial diversity in *Porites lobata* between the wet and dry seasons. The initial intention of the project was to sample Inner and Outer sites in Fouha Bay for two four-month periods that transitioned from the wet to the dry season. However, the wet season ended earlier than anticipated nearing the end of November 2019 that led to samples being collected during a 3-month period of the wet season and a 5-month period of the dry season (Figure 1). It is worth noting that there were two typhoons, Typhoon Krosa and Typhoon Francisco (Wikipedia contributors, 2026), that originated near Guam at the beginning of August 2019 that delayed the start of this project and led to high precipitation and runoff. These severe tropical disturbances led to a large influx of sediment into Fouha Bay that likely led to the inundation of both inner and outer site with sedimentation similar to what has been observed during the wet season previously (Lock et al., 2024). This likely influenced the microbial diversity at the first time point in the study (i.e., increased bacterial diversity in September at the outer site; Fig. 5).

### Symbiodiniaceae Diversity

Most species of *Porites* have been thought to be associated with *Cladocopium* C15 (Lajeunesse et al., 2004; Thornhill et al., 2006). However, more recently massive *Porites* species were described to harbor mixed communities of *Cladocopium* and *Durusdinium* (Terrano et al., 2019; Tan et al., 2020). Species of *Porites* generally transmit their Symbiodiniaceae endosymbionts vertically (Bennett et al., 2024), a likely explanation of the relatively low variability of Symbiodiniaceae observed in this study. ITS2 metabarcoding indicated that *P. lobata* samples were dominated by *Cladocopium* C15 and several of its variants (Fig. 3), a result consistent with prior studies that identified *Cladocopium* C15 and its variants to dominate *P. lobata* in Guam (Lock et al., 2025) and *Porites* in general (e.g., Hoadley et al., 2021; Grupstra et al., 2024). Nonetheless, inner and outer site Symbiodiniaceae community composition was significantly different (Table 1), a likely result of variable contributions of *Cladocopium* C15 variants between sites (Fig. 3; Supplementary Fig. 1).

A Pearson χ² test employed in a previous study (Fifer et al., 2022) indicated that Symbiodiniaceae ITS2 type profile frequencies did not differ significantly among sites, suggesting that dominant symbiont identities were broadly consistent across sites. However, the multivariate analysis of ITS2 variants employed herein (PERMANOVA) revealed significant differences in Symbiodiniaceae community structure among sites (Table 1), indicating site-specific variation in background symbiont assemblages. Taken together, these results indicate that the dominant Symbiodiniaceae identity of *P. lobata* was conserved across sites in Fouha Bay, but background ITS2 variant compositions differ. This may suggest host filtering of dominant C15 Symbiodiniaceae symbionts paired with environmental structuring of background variants. Differences in *Cladocopium* C15 variant functional traits have previously been identified and linked to differences in *Porites* stress resilience (Hoadley et al., 2021). It is conceivable that *Cladocopium* C15 variants variation in *P. lobata* in Fouha Bay is linked to habitat differences, but further study of Symbiodiniaceae traits will be necessary to shed light on this possibility.

It is noteworthy that some *P. lobata* samples contained *Gerakladium* G3x (Fig. 3), a member of a relatively poorly studied Symbiodiniaceae genus. While *Gerakladium* is typically associated with sponges and soft corals (van Oppen et al. 2005; Pochon et al. 2007; Granados et al. 2008; Hill et al. 2011), a study conducted in Hawai’i found *Gerakladium* in *P. lobata* and concluded that this genus may be horizontally transferred from bio-eroding sponges to the coral colony (Stat et al., 2013). *Gerakladium* is thought to contain thermotolerant variants of Symbiodiniaceae that are found in different host corals growing in suboptimal conditions (Mote et al., 2021), which may explain *Gerakladium*’s presence in Fouha Bay. However, *Gerakladium* relative abundances did not appear to show any clear patterns across samples, sites, or seasons (Fig. 3), and were found in relatively low abundances when observed. Given these considerations, it is also possible that *Gerakladium* presence is the result of stochastic communities that live on the surface or in the gastrovascular cavity of *P. lobata* rather than representing endosymbionts (LaJeunesse et al., 2018). Further study will be necessary to better understand *Gerakladium*’s association with *P. lobata*.

### Bacterial Microbiome Diversity

A previous study of *P. lobata* microbiomes along the Fouha Bay sedimentation gradient identified *Endozoicomonadaceae* as the dominant taxon of *P. lobata* microbiome communities (Fifer et al., 2022), consistent with the results presented here (Fig. 6). This dominance of *Endozoicomonadaceae* has been shown in *P. lobata* and other coral species sampled across different reefs in Guam (Lock et al., 2025; Miller & Bentlage, 2024). These findings from Guam are consistent with broader patterns of coral microbiome diversity across the Pacific (Galand et al., 2023). *Endozoicomonadaceae* are generally considered beneficial to corals (e.g., Neave et al., 2016), especially when exposed to stressful conditions (e.g., Maher et al., 2020), but their role as purely beneficial endosymbionts has recently been questioned (Pogoreutz & Ziegler, 2024). In this context, it is worth noting that *Porites* in Guam appears to host species of *Parendozoicomonas* (Lock et al., 2025) similar to *Porites* spp. sampled across the broader Pacific Ocean (Hochart et al., 2023). Most of our knowledge of the coral-*Endozoicomonadaceae* relationship comes from genome-enabled bioinformatic inferences of *Endozoicomonas* spp. traits while *Parendozoicomonas* information of, for example, its metabolic repertoire remains severely limited at this point.

Previous sampling of *P. lobata* from a single time point during the dry season found no significant difference in microbiomes along the Fouha Bay sedimentation gradient (Fifer et al., 2022). By contrast, the time series data presented here found that bacterial diversity differed significantly between sites and seasons, with the interaction term of site/season contributing to differentiation between samples (PERMANOVA: site R² = 0.11, season R² = 0.05, interaction R² = 0.03; Table 2). While site, season, and their interaction were significant in explaining bacterial community variation, within site variability was high (residual R² = 0.81). High residual variability is common in coral microbiome studies, reflecting substantial colony-level heterogeneity and the presence of a variable microbial community of rare and transient taxa beyond a small core microbiome (e.g., Ainsworth et al., 2015; Hernández-Agreda et al. 2018; Hernández-Agreda et al. 2018; Morrow et al. 2022). Given these considerations, it is likely that spatial structuring of bacterial communities was apparent in our dataset because temporal community variation was incorporated in the analysis, highlighting the need for further work emphasizing seasonal sampling designs.

Site was likely the largest driver of microbial diversity due to the inner site’s proximity to the river mouth, close to freshwater runoff and higher accumulation of sediments compared to the outer site (Lock et al., 2024). The highest bacterial diversity was found at the inner site during the wet season, a result that was also reflected in increased relative abundances across bacterial groups at the inner site during the wet season compared to the dry season and the outer site (Fig. 6). Elevated diversity and high relative abundance of bacterial groups was a likely result of the inner site’s proximity to the river mouth combined with increased runoff, sedimentation, and available nutrients during the wet season. Increased bacterial diversity was likely caused by chronic nutrient and sediment availability that increased influx of microbes, with coral microbiomes shifting to facilitate persistence of the host under harsh environmental conditions (Rosenberg et al., 2007). The high sediment load experienced at the inner site during the wet season may have allowed rarer taxa to outcompete *Endozoicomonadaceae,* which decreased from 82.5% relative abundance across all samples to 71.7% during this time period. A caveat to this interpretation is that the data presented here represent relative abundances; *Endozoicomonadaceae* abundance could have remained constant while overall abundance of bacteria in the microbiome could have increased. By contrast, microbiomes at the outer site were highly stable across sampling time points and seasons, likely due to similar environmental conditions between the wet and dry seasons as well as increased flushing rates that occur with further distance from the mouth of the river. Interestingly, massive *Porites* sampled from extreme reef sites (high temperature and light attenuation) in Palau have been described as harboring few *Endozoicomonadaceae*, and in this case microbiome differences across sites were linked to host coral lineage (Grupstra et al., 2024). While differences in bacterial microbiomes observed between inner and outer sites in Fouha Bay could be related to *P. lobata* lineage identity, a previous study assigned the majority (>90%) of *P. lobata* colonies sampled from the area of Fouha Bay that encompasses our inner and outer sites to a single lineage (clade V; Primov et al., 2024).

*P. lobata* in Fouha Bay was dominated by *Endozoicomonadaceae* across all sites and seasons, but additional taxa were uniquely affected by environmental conditions. While the microbiomes across all *P. lobata* colonies were largely dominated by two bacterial families, *Endozoicomonadaceae* and *Oxalobacteraceae* (mean relative abundance of 87.4% and 4.7%), each site/season group was characterized by additional bacterial groups. The inner site during the dry season was characterized by a higher relative abundance of Endozoicomonadaceae and an increase in Oxalobacteraceae, particularly members of the genus *Herbaspirillum*. Species within *Herbaspirillum* are commonly associated with terrestrial plants and soils, including as endophytic diazotrophs (Schmid et al., 2006; Monteiro et al., 2012), suggesting a potential allochthonous origin. The observed enrichment may be consistent with inputs of terrestrially derived material and suspended particles, as well as reduced flushing and increased residence times that can facilitate the retention of particle-associated microbial taxa in nearshore environments (Crump et al., 2004; Fortunato and Crump, 2015; Dang and Lovell, 2016). The inner site wet season group was characterized by *Endozoicomonadaceae* and showed an increase in other rare taxa, likely a result of severe sedimentation in the proximity of the La Sa Fua River (Minton et al., 2022; Lock et al., 2024). It is worth noting that there were multiple tropical disturbances during this study that led to increased sedimentation and resuspension rates that may have increased microbial diversity, although long-term data would need to be collected to gain further insight into longer-term trends of bacterial microbiome diversity close to the mouth of Fouha Bay.

Outer site dry season samples showed the highest microbial community stability and lowest diversity, a likely result of the overall stable and more favorable environmental conditions compared to the inner site and wet season. By contrast, the inner site wet season sample group experienced the harshest environmental conditions that led to low microbiome stability (changes in taxonomic composition compared to the dry season) and highest bacterial diversity among samples collected. Interestingly, outer site wet season samples saw an increase in *Vibrionaceae* compared to the remainder of sample groups*. Vibrionaceae* are heterotrophic bacteria that dominate in coastal areas where they contribute to organic carbon cycling and may rapidly proliferate in response to nutrient inputs (Zhang et al., 2018). Their high abundance at the outer site during the wet season may have been the result of dissolved sediment load and nutrients that were mixing with forereef seawater towards the outside of Fouha Bay.

The four unique site and season groups highlight the microbial flexibility that exists within *P. lobata* microbiomes. Typically, *Acropora* species have been termed “microbiome conformers’’ that experience shifts in their microbial diversity due to environmental conditions while corals like *Pocillipora* species have been termed “microbiome regulators” that maintain a constant microbiome despite environmental shifts (Ziegler, 2019). Past studies have shown that *Porites* may act as a microbiome regulator rather than a microbiome conformer (Maher, 2019). Our results from Fouha Bay show that the microbiome of *P. lobata* was dominated by *Endozoicomonadaceae* across sites and seasons but differences in diversity and microbiome composition were apparent between sites and seasons. These differences in diversity were likely a result of environmental differences, with high precipitation and resulting sedimentation increasing diversity and taxonomic composition of bacterial microbiomes at the inner site during the wet season most drastically. Indeed, bacterial groups that may be of terrestrial origin (Actynomycetota and Deinoccoccota) were over-represented at the inner site and/or during the wet season. Deinococcales have been described from disturbed soils (e.g., Rainey et al., 2005), such as those found in the upstream badlands of the Fouha watershed. Actinomycetes have been identified as constituents of coral microbiomes previously, with the potential to produce bioactive metabolites (e.g, Mahmoud & Kalendar, 2016) and to increase coral resilience (e.g., Li et al., 2023). However, actinomycetes are particularly well-known from soils where they represent a major reservoir of antibacterial genes and antibiotic resistance genes (ARGs; McClung et al., 2022). Their elevated abundance in near-shore marine ecosystems, such as Fouha Bay, warrants further study given the potential environmental health risks associated with the proliferation of ARGs (e.g., Muteeb et al., 2025). Elevated abundance of Clostridia during the wet season may be associated with low oxygen conditions caused by increased sedimentation. Indeed, such increases in Clostridia have been associated with the deoxygenation of seawater which favors this group of facultative anaerobes (Howard et al., 2023). While Clostridia are known to cause disease in humans and animals (Cruz et al., 2019), their role in coral microbiomes remains ambiguous (Howard et al., 2023).

Given the results presented here that highlight differences in *P. lobata* microbiomes across Fouha Bay, longer-term monitoring should further investigate the dynamic nature of *P. lobata* microbiomes and identify patterns of microbial recovery following wet season disturbances (Ziegler, 2019). Considering the resilience of *P. lobata*, a coral that persists under high sedimentation stress, further insights into the ability of individual coral colonies to revert back to a stable microbiome when chronic stress conditions subside would be valuable. In addition to differences in bacterial microbiome diversity across sites and seasons, we found a weak but positive correlation between bacterial and Symbiodiniaceae diversity (Supplementary Fig. 2) that may be indicative of Symbiodiniaceae and bacterial communities associated with *P. lobata* to be affected in a similar fashion by habitat differences. Further monitoring and understanding of the correlated community dynamics of bacteria and Symbiodiniaceae across seasons would provide valuable insights into the connections between these two constituents of the coral holobiont.

## ACKNOWLEDGEMENTS

This work was supported by Guam NSF EPSCoR through the National Science Foundation (awards OIA-1457769 and OIA--1946352).

## SUPPLEMENTARY TABLES

**Supplementary Table 1.** ANOVA table showing the contributions of site, season, and their interaction term on observed diversity of Symbiodiniaceae communities.

|  | *df* | *Sums of Squares* | *Mean Square* | *F* | *p* |
| --- | --- | --- | --- | --- | --- |
| **Site** | 1 | 1.15E+10 | 1.15E+10 | 426.791 | <2e-16 *** |
| **Season** | 1 | 8.45E+05 | 8.45E+05 | 0.031 | 0.86 |
| **Site:Season** | 1 | 4.64E+07 | 4.64E+07 | 1.728 | 0.191 |
| **Residuals** | 108 | 2.90E+09 | 2.69E+07 |  |  |

**Supplementary Table 2.** ANOVA table showing the contributions of site, season, and their interaction term on evenness of Symbiodiniaceae communities.

|  | *df* | *Sums of Squares* | *Mean Square* | *F* | *p* |
| --- | --- | --- | --- | --- | --- |
| **Site** | 1 | 0.01283 | 0.012835 | 31.5 | 1.56e-07 *** |
| **Season** | 1 | 0.00087 | 0.000867 | 2.128 | 0.147 |
| **Site:Season** | 1 | 0.00004 | 0.000036 | 0.09 | 0.765 |
| **Residuals** | 108 | 0.044 | 0.000407 |  |  |

**Supplementary Table 3.**
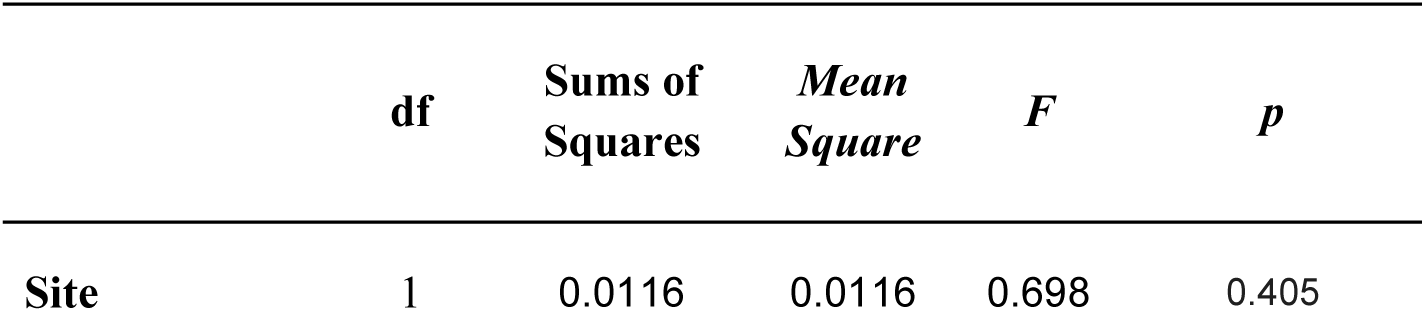
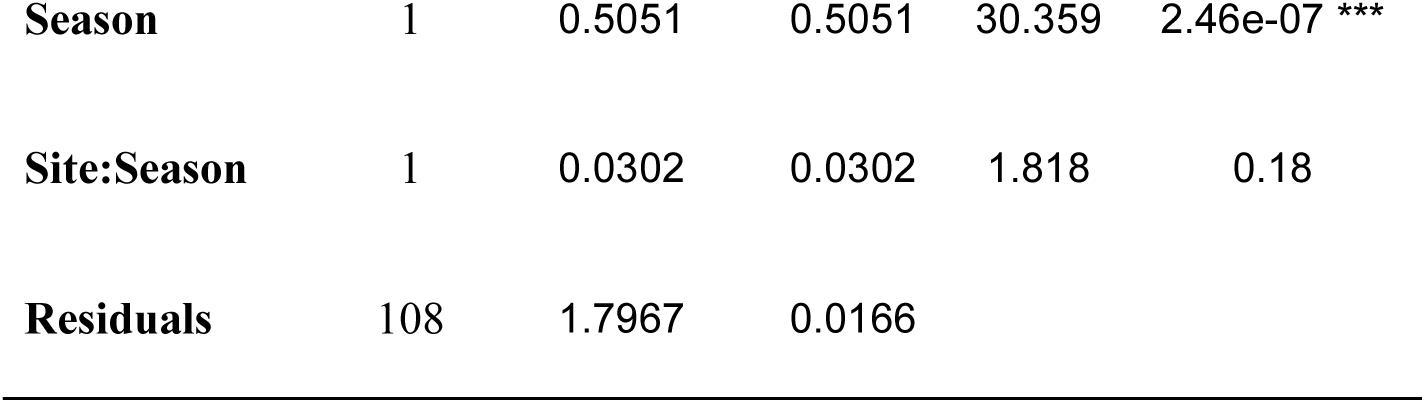
ANOVA table showing the contributions of site, season, and their interaction term on Shannon diversity of Symbiodiniaceae communities.

**Supplementary Table 4.** ANOVA table showing the contributions of site, season, and their interaction term on observed diversity of bacterial communities.

|  | **df** | **Sums of Squares** | **Mean Square** | **F** | **p** |
| --- | --- | --- | --- | --- | --- |
| **Site** | 1 | 65714 | 65714 | 8.13 | 0.00522 ** |
| **Season** | 1 | 205631 | 205631 | 25.44 | 1.85e-06 *** |
| **Site:Season** | 1 | 244984 | 244984 | 30.31 | 2.51e-07 *** |
| **Residuals** | 108 | 872910 | 8082 |  |  |

**Supplementary Table 5.** ANOVA table showing the contributions of site, season, and their interaction term on evenness of bacterial communities.

|  | **df** | **Sums of Squares** | **Mean Square** | **F** | **p** |
| --- | --- | --- | --- | --- | --- |
| **Site** | 1 | 0.2065 | 0.20653 | 22.067 | 7.79e-06 *** |
| **Season** | 1 | 0.2091 | 0.20912 | 22.343 | 6.92e-06 *** |
| **Site:Season** | 1 | 0.0319 | 0.0319 | 3.409 | 0.0676 |
| **Residuals** | 108 | 1.0108 | 0.00936 |  |  |

**Supplementary Table 6.**
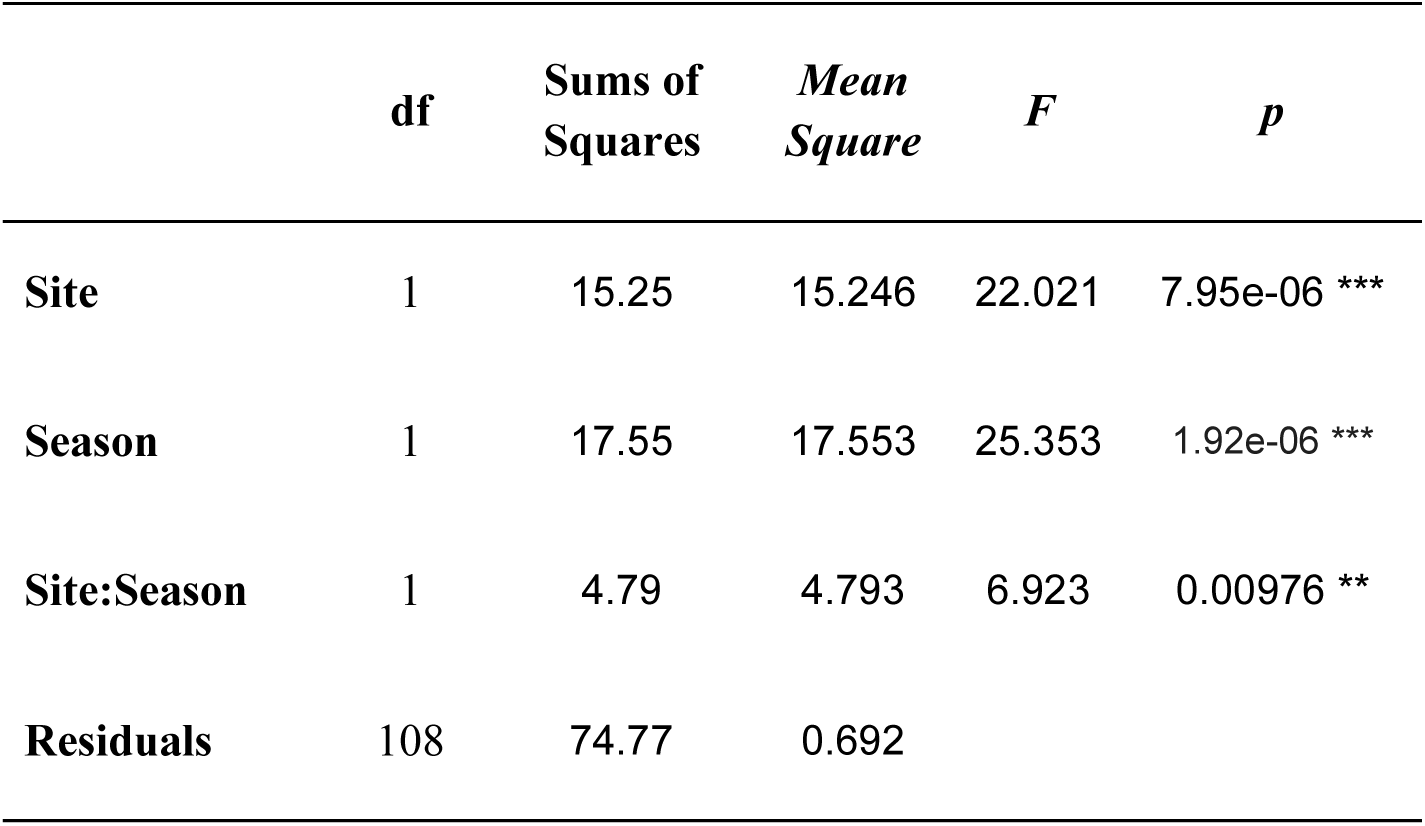
ANOVA table showing the contributions of site, season, and their interaction term on Shannon diversity of bacterial communities.

|  | **df** | **Sums of**<br>**Squares** | ***Mean***<br>***Square*** | ***F*** | ***p*** |
| --- | --- | --- | --- | --- | --- |
| **Site** | 1 | 15.25 | 15.246 | 22.021 | 7.95e-06 *** |
| **Season** | 1 | 17.55 | 17.553 | 25.353 | 1.92e-06 *** |
| **Site:Season** | 1 | 4.79 | 4.793 | 6.923 | 0.00976 ** |
| **Residuals** | 108 | 74.77 | 0.692 |  |  |

## SUPPLEMENTARY FIGURES

**Supplementary Fig. 1.**
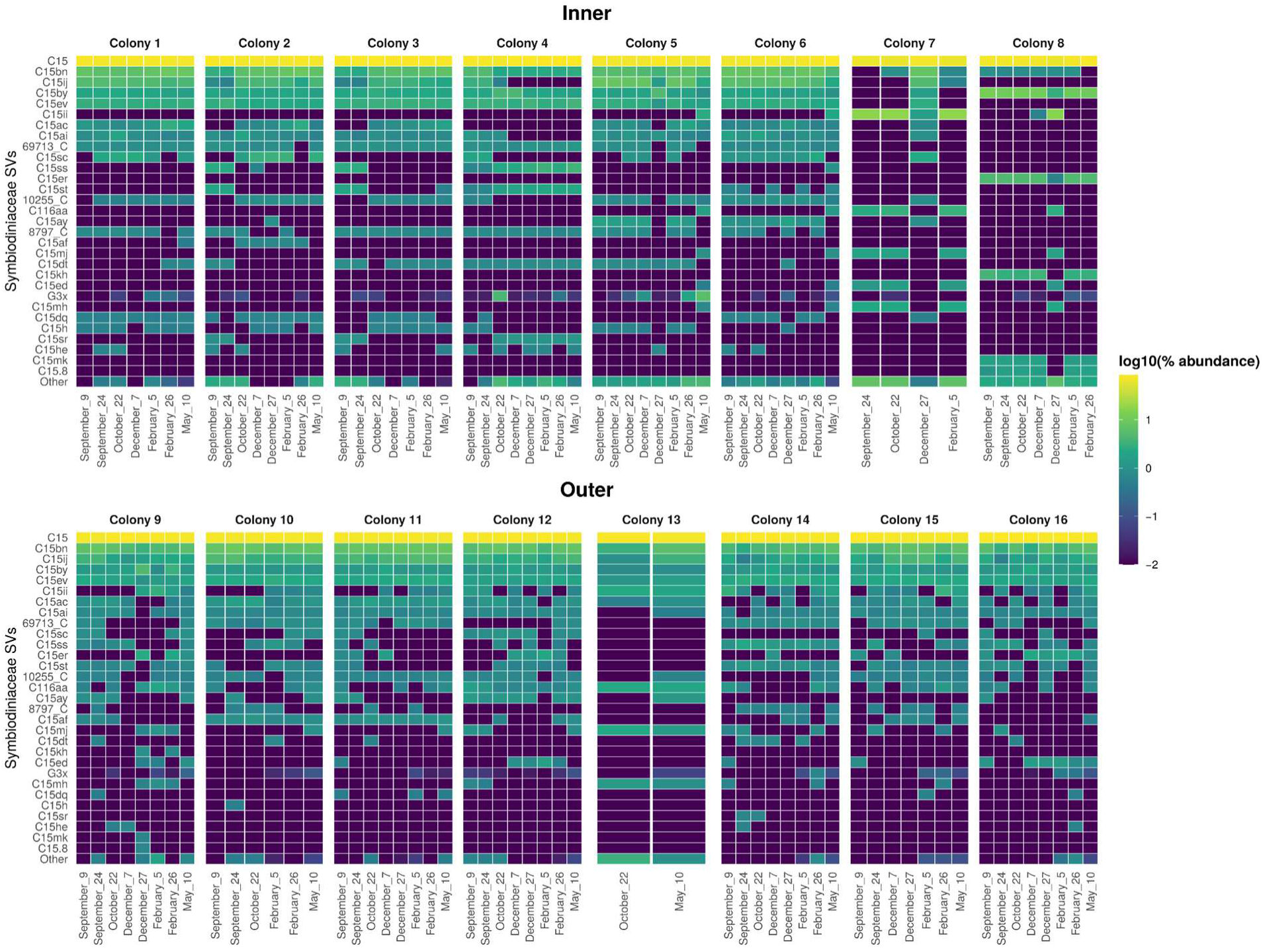
Symbiodiniaceae community composition at inner and outer sites for each sampled colony. Columns of the heat map represent individual samples; rows represent Symbiodiniaceae clades. Dates samples were taken are denoted on the horizontal axis. Relative abundances were log-scaled using the common logarithm where 2 equals 100% (yellow) and -2 equals 0.01% (blue). Symbiodiniaceae clade abbreviations: C = *Cladocopium*; G = *Gerakladium*. SV = sequence variant.

**Supplementary Fig. 2.**
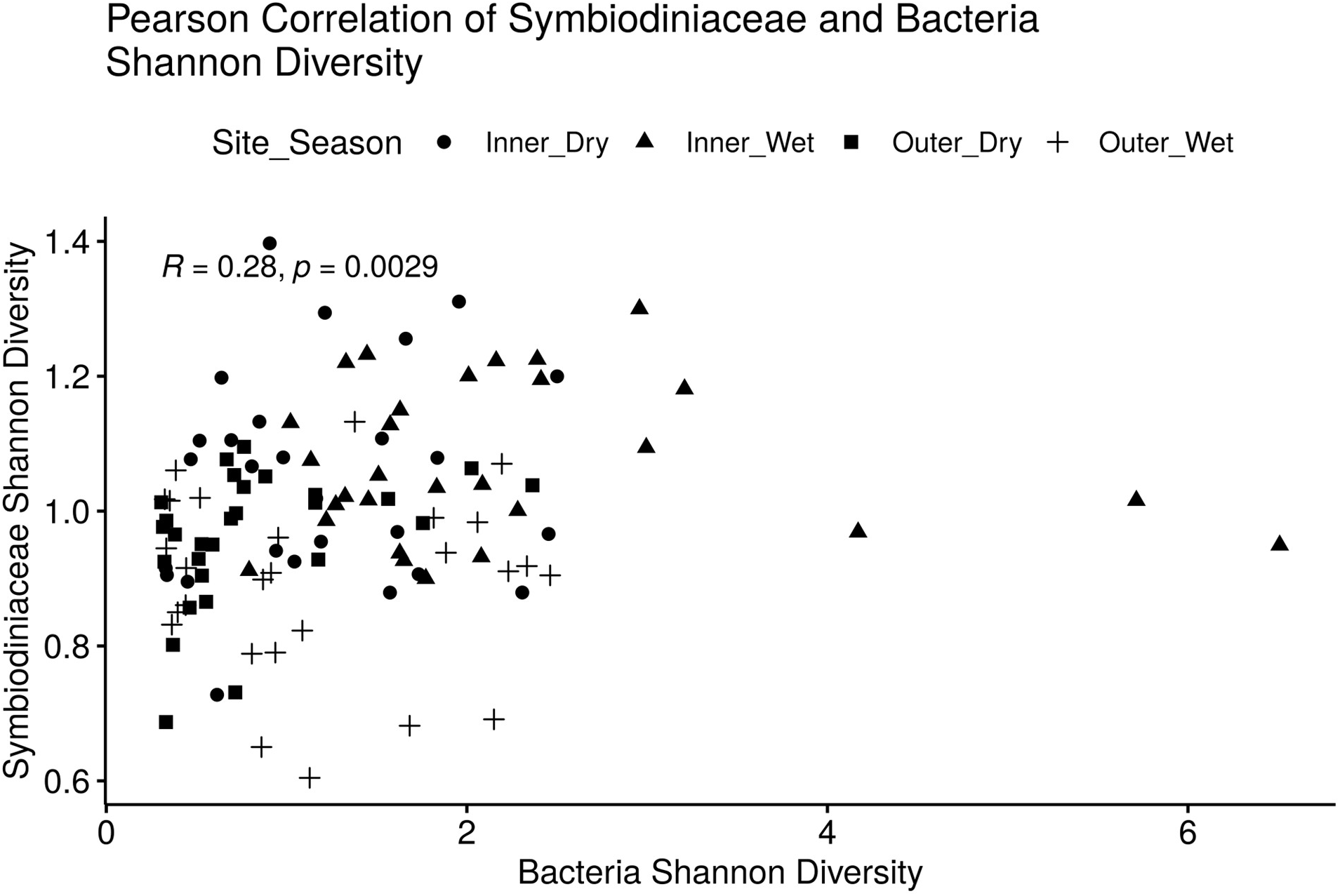
Pearson correlation of Shannon diversity of Symbiodiniaceae and bacterial communities. Samples were grouped by site (inner vs. outer bay) and season (dry vs. wet).

**Supplementary Fig. 3.**
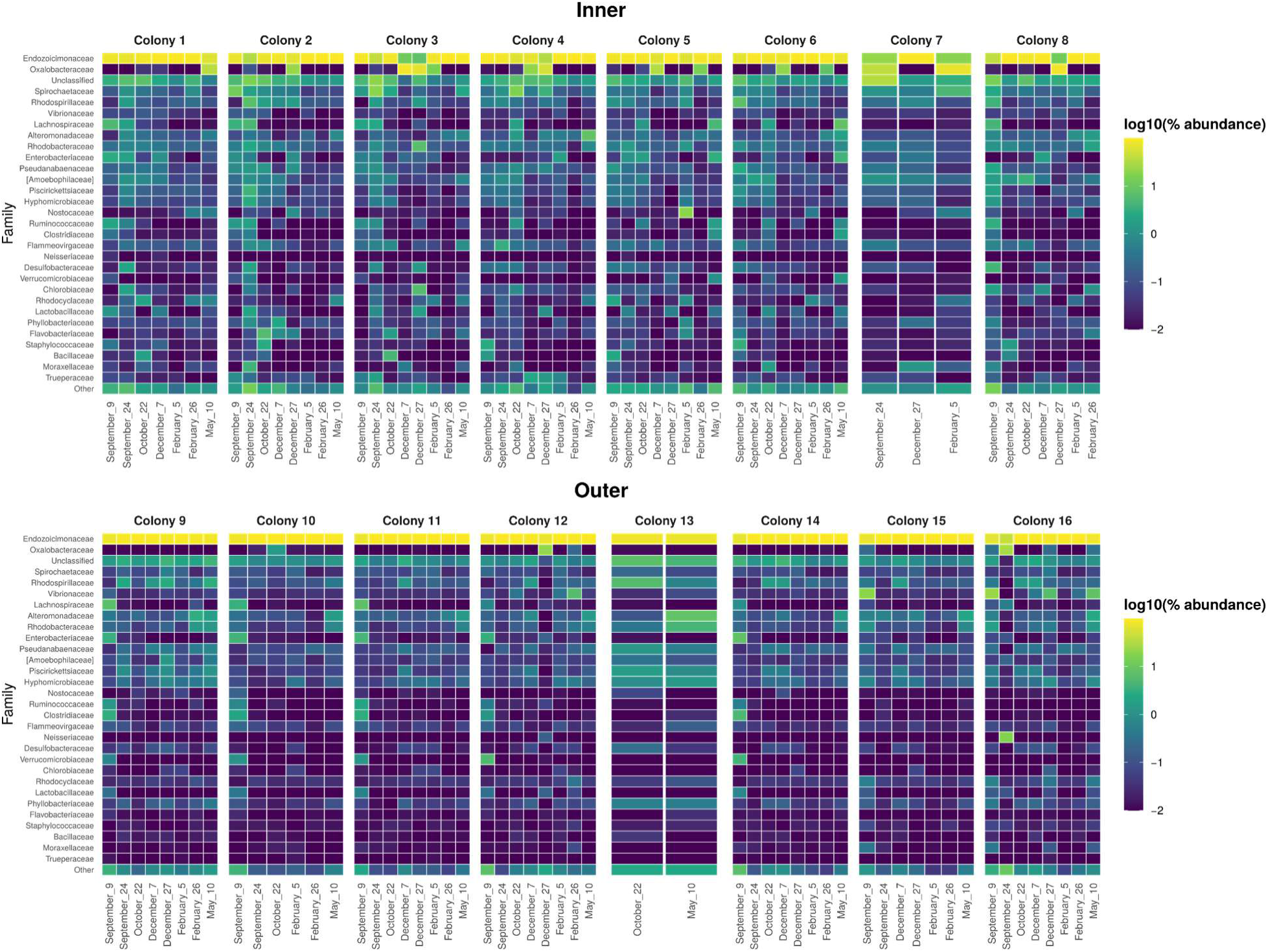
Relative abundance of the 30 most abundant bacterial families in *Porites lobata* microbiomes sampled from inner and outer site in Fouha Bay. Columns of the heat map represent samples taken from each colony, with sampling dates denoted along the horizontal axis; rows represent bacterial taxa inferred from ASVs. Relative abundance was log-scaled using the common logarithm where 2 equals 100% (yellow) and -2 equals 0.01% (blue).

